# Validation and Application of a Bench Top Cell Sorter in a BSL-3 Containment Setting

**DOI:** 10.1101/2020.07.30.229146

**Authors:** Lydia M Roberts, Rebecca Anderson, Aaron Carmody, Catharine M Bosio

## Abstract

Rigorous assessment of the cellular and molecular changes during infection typically requires isolation of specific immune cell subsets for downstream application. While there are numerous options for enrichment/isolation of cells from tissues, fluorescent activated cell sorting (FACS) is accepted as a method that results in superior purification of a wide variety of cell types. Flow cytometry requires extensive fluidics and aerosol droplets can be generated during collection of target cells. Pathogens such as *Francisella tularensis*, *Mycobacterium tuberculosis*, *Yersinia pestis*, and SARS-CoV-2 require manipulation at biosafety level-3 (BSL-3). Due to the concern of potential aerosolization of these pathogens, use of flow cytometric-based cell sorting in these laboratory settings requires placement of the equipment in dedicated biosafety cabinets within the BSL-3. For many researchers, this is often not possible due to expense, space, or expertise available. Here we describe the safety validation and utility of a completely closed cell sorter that results in gentle, rapid, high purity, and safe sorting of cells on the benchtop at BSL-3. We also provide data demonstrating the need for cell sorting versus bead purification and the applicability of this technology for BSL-3 and potentially BSL-4 related infectious disease projects. Adoption of this technology will significantly expand our ability to uncover important features of the most dangerous infectious diseases leading to faster development of novel vaccines and therapeutics.

## Introduction

The ability to sort pure populations of immune cells is a critical tool for immunologists. Traditionally, multi-parameter, droplet-based cell sorters have been used to isolate cells for downstream analyses. The utility of these instruments is undeniable, and they have allowed for a plethora of important advances in a wide variety of fields. However, they have inherent limitations. First, there is a level of cell loss that is not easily, if at all, controlled. This can require the use of additional animals to compensate for loss. Second, depending on the size of the target cell population, the time required to collect that population can be limiting for additional analyses. Lastly, there is the propensity for the generation of aerosols during the sorting process.

Due to the potential of aerosol exposure to the sort operator, a risk assessment should be performed prior to working with unfixed samples. Fontes, et al provides the current International Society for the Advancement of Cytometry guidelines for sorting samples at all biosafety levels [1]. There are a number of high consequence pathogens such as *Francisella tularensis*, *Mycobacterium tuberculosis*, *Yersinia pestis*, and SARS-CoV-2, that require manipulation at biosafety level 3 (BSL-3). Handling these organisms at BSL-3 is necessary because one of the primary risks of these pathogens is the ability to cause pulmonary infection following inhalation of aerosols. The risk of aerosolization of infectious material during sorting has been partially mitigated with built-in engineering controls such as the Aerosol Management System for the FACSAria II and the placement of cell sorters in biosafety cabinets (BSC) [2]. However, the potential to generate aerosols in combination with the aforementioned challenges remains an insurmountable hurdle for many laboratories in need of sorting cells to a high degree of purity.

Ideally, cell sorting in the BSL-3 laboratory setting should fulfill the following criteria. First, the process should be safe with limited or the complete absence of aerosol generation. Second, the sorting process should be efficient with a reduction or cessation of cell loss. Third, there is the capability to sort using multiple parameters to increase identification of targeted cell populations. Fourth, the sample should remain sterile. Finally, the equipment must be straightforward to use and maintain to facilitate the routine application of cell sorting for immunological studies. Recently, Miltenyi Biotec unveiled a completely closed cell sorting system they coined the MACSQuant Tyto (Tyto). This system fulfills all of the requirements described above for potential use in containment settings. However, prior to application it was necessary to validate the absence of the generation of aerosol following a sort. Further, it was also necessary to confirm the activity of sorted populations from the Tyto.

Herein, we describe validation procedures for use of the Tyto in BSL-3 settings. Additionally, we provide a quality control method to periodically check instrument performance. We also include data demonstrating that cell sorting using this technology resulted in not only a more rapid procurement of cell populations with decreased hands on time by the user, but also resulted in the identification of important contributions of low numbers (<15%) of contaminating cells to distinct elements of cellular metabolic activity among cells isolated from the lung.

## Materials and Methods

### Aerosol testing

Internally fluorescent (excitation 480 nm; emission 520 nm) 1.0 μm Dragon Green beads (Bangs Laboratories, Inc.) were diluted 1:100 in Tyto running buffer (Miltenyi Biotec). Aerosol testing was performed on a FACSAria II (BD Biosciences) with the Aerosol Management System disabled or a MACSQuant Tyto (Tyto; Miltenyi Biotec). Aerosol samples were collected using a Cyclex-d impactor sampling cassette, MegaLite pump, and Rotameter (Environmental Monitoring Systems) using a constant vacuum set to 20 l/minute. Samples were collected for up to 30 seconds on the FACSAria II, 10 minutes with the Tyto door closed, and 30 seconds with the Tyto door open immediately after the sort completed. Following sample collection, the coverslip was inverted onto a microscope slide and viewed using an Axio Imager (Zeiss).

### Quality control testing

Veri-cells (Biolegend) were reconstituted according to the manufacturer’s instructions in 2 ml of Tyto running buffer and then loaded into the sort cartridge. CD4^+^ cells were sorted on the Tyto using PE as the trigger channel and the VioBlue channel to determine cell speed. Input, sorted, and negative fractions were analyzed on the Symphony flow cytometer (BD Biosciences) and subsequently in FlowJo 10 (BD Biosciences). Singlets were gated by plotting FSC-H versus FSC-A. From the singlet gate, cells were gated by plotting SSC-A versus FSC-A. Within the cell gate, CD4+ cells were gated by plotting CD3 Pacific Blue versus CD4 APC. Sort efficiency was calculated as follows: (number of target cells in the sort fraction) / (number of target cells in the sort fraction) + (number of target cells in the negative fraction). Depletion yield was calculated as follows: (percentage of target cells in input fraction – percentage of target cells in the negative fraction) / (percentage of target cells in the input fraction).

### Mice

Five- to seven-week-old female C57Bl/6J mice were purchased from Jackson Laboratories and housed at ABSL-2 at Rocky Mountain Laboratories. Prior to euthanasia, circulating T cells were intravenously labeled for 3 minutes via injection of 2.5 μg anti-CD45.2 FITC in 100 μl of sterile saline as previously described [3]. All animal studies were approved by and conducted in accordance with RML’s Animal Care and Use Committee.

### CD4^+^ T cell purification

Lungs were aseptically removed and digested into a single cell suspension as previously described [4]. CD4^+^ T cells were enriched from all samples using a CD4 TIL Microbeads kit (Miltenyi Biotec) according to the manufacturer’s instructions. The resulting enriched cells were then stained with anti-CD4 BV421 (clone GK1.5), anti-CD44 PE-Cy7 (clone IM7), and anti-Thy1.2 APC (clone 30-H12) (Biolegend). Total circulating CD4^+^ T cells were sorted on the Tyto using CD4 BV421 as the cell trigger and CD45.2 FITC as the cell speed channel. All samples were >98% pure as determined by flow cytometry analysis.

### Metabolic flux analysis

All T cell activation assays to determine changes in metabolic flux were performed as previously described [5]. Briefly, 2×10^5^ purified T cells were seeded per well in a poly-D-lysine (100 μg/ml) coated Seahorse 96 well plate and centrifuged for 5 minutes at 300xg. Cells were incubated for 1 hour in a non-CO_2_, 37°C incubator prior to analysis. After 4 baseline measurements, T cells were activated by injection of anti-CD3/CD28 beads (Miltenyi Biotec) at a ratio of 8:1 beads:T cell followed by injection of oligomycin (2μM) and then 2-deoxyglucose (2-DG; 50 mM). The extracellular acidification rate (ECAR) was measured using a Seahorse XFe96 Bioanalyzer (Agilent) with readings collected approximately every 6.5 minutes.

### Statistics

Statistically significant differences between two groups was determined using an unpaired, two-tailed t-test. Significance was set at *p* ≤ 0.05.

## Results

### Absence of Detectable Generation of Aerosols During Use of the Tyto

As described above the generation of aerosols during cell sorting procedures is an important impedance to operation of flow cytometric based cell sorting in containment settings. Therefore, prior to usage of the Tyto on the benchtop to sort cells at BSL-3, it was necessary to empirically determine if aerosols were generated during the cell sort procedure. We utilized a novel method described by Perfetto, et al [6] using internally fluorescent 1.0 μm beads to perform aerosol testing on the Tyto. These beads were uniform in size and intensely fluorescent in the FITC channel (figure 1A). As a positive control, we detected aerosols generated by the in-house FACSAria II when the Aerosol Management System was disabled, and the flow stream disrupted. As expected, beads were detected on the coverslip after only a 30 second exposure (figure 1B).

**Figure 1.**
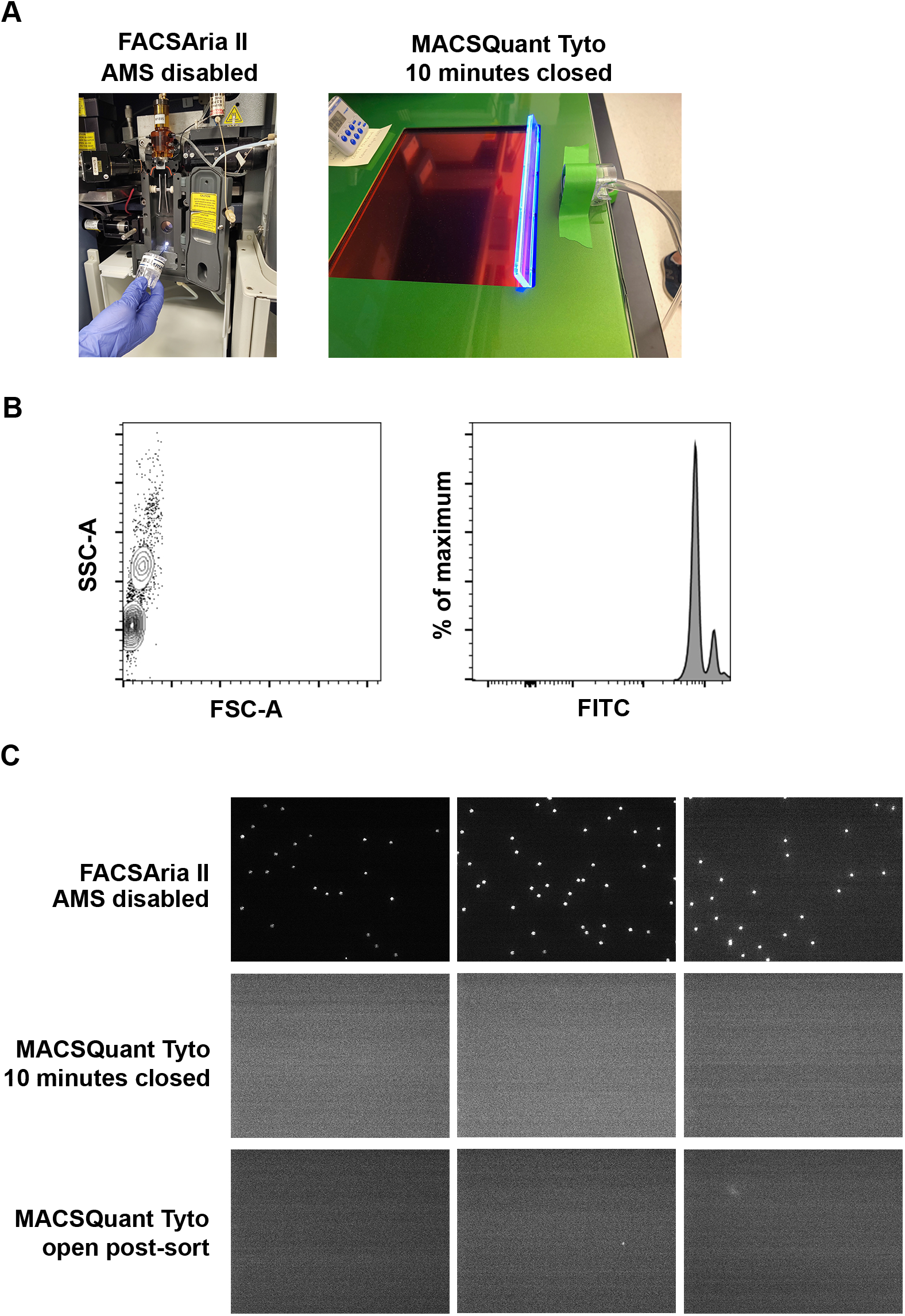
Aerosols are not generated by the Tyto cell sorter. A) Aerosol testing impactor set-up on the FACSAriaII with AMS disabled and the Tyto. B) A solution containing 1.0 μm internally fluorescent beads in the FITC channel were used for aerosol testing. C) Representative images of coverslips analyzed after aerosol testing on the FACSAria II or Tyto with a closed (during sort) or open (post-sort) door. Aerosol testing was performed in triplicate for each condition. The entire coverslip from each Tyto test was scanned and no beads were detected.

During aerosol testing of the Tyto, we recapitulated instrument use during a cell sort. The manufacturer’s instructions were followed to prime and load the sort cartridge into the instrument. The Tyto will only sort if the door is closed, thus the impactor cassette was placed adjacent to the closed door and allowed to run for 10 minutes while the fluorescent beads were sorted. If aerosols were generated during the sort procedure, the user could be exposed when the Tyto’s door is opened at the sort’s completion for removal of the cartridge. Therefore, at the end of the 10 minutes, the impactor cassette was removed and replaced with a new one prior to opening the Tyto door. The new impactor cassette was used to sample next to the sort cartridge while it was still in the instrument. We sampled the entire duration the door remained open before it defaulted to closing; the sample time was approximately 30 seconds. All coverslips were visualized using the same gain and focal plane where beads were detected from our positive control. We did observe autofluorescent debris which clearly differed in size and fluorescent intensity compared to the beads. This observation wasn’t surprising given the amount of air that was passed across the coverslip during the testing procedure. The entire coverslip was scanned; no beads were detected from samples collected when the Tyto door was open or closed (figure 1b). These data indicate that aerosols were not generated during the cell sort procedure and the Tyto can safely be used on the benchtop at BSL-3.

### Quality control testing of Tyto

Routine quality control testing of laboratory equipment ensures consistent instrument performance and can reveal mechanical issues prior to their use on precious samples. Standard testing procedures also allows for new users to be trained on the equipment with an expected and historically consistent outcome. To this end, we established a quality control sort to periodically verify the Tyto was performing as expected. We selected lyophilized Veri-cells as a commercially available, consistent sample to sort cells from using a standardized sample volume, cell concentration, and gating strategy. CD4^+^ T cells were approximately 20% of the input sample and were sorted to 97% purity (figure 2A). Because there is no sample loss when sorting on the Tyto, the negative fraction could also be analyzed to determine the extent of target population depletion (figure 2A). The depletion yield was >80% and the calculated sort efficiency was >85% (figure 2B). This quality control analysis was performed multiple times over the six-plus months the instrument has been in use and demonstrate the stability of the Tyto over time in our hands. Furthermore, at least one month passed between each quality control analysis and it was often the case the machine was not utilized during this time. Our laboratory has established a purity of >96%, depletion yield of >80%, and sort efficiency of >85% as benchmarks that must be met during quality control testing.

**Figure 2.**
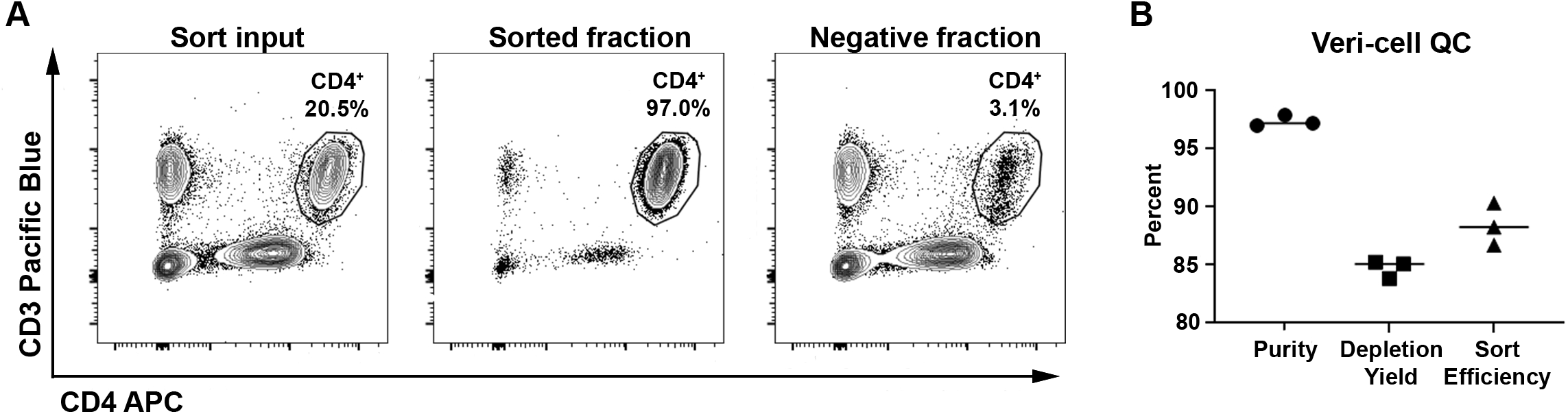
Establishing quality control sort of Veri-cells. A) Representative flow plots for the Veri-cell sort input, sorted fraction, and negative fraction showing the percentage of CD4^+^ T cells in each sample analyzed on a BD Symphony flow cytometer using the gating strategy described in the Material and Methods. B) The percent purity, depletion yield, and sort efficiency of 3 independent quality control sorts was calculated.

### Tyto sorted cells are highly pure and superior to bead purified cells

We previously established that anatomical location within the pulmonary compartment influences a CD4^+^ effector T cell’s (T_eff_) glycolytic capacity [5]. As an internal control for this study, we included total pulmonary CD4^+^ T cells isolated from naïve mice in each metabolic flux experiment. Due to unavoidable time constraints in sorting resident and circulating pulmonary T cells from immune animals, it was not logistically possible to also sort total CD4^+^ T cells using the FACSAria II from naïve mice. Rather, total pulmonary CD4^+^ T cells were purified from naïve mice using CD4 microbeads. When analyzed for metabolic flux, cells from naive mice routinely presented with an elevated basal extracellular acidification rate (ECAR) compared to our sorted immune controls. Post-purification flow analysis revealed total naïve CD4^+^ T cells consistently had 10-15% of contaminating lung cells. Therefore, we attributed the elevated level of basal ECAR in our naïve CD4^+^ T cell samples to these contaminating cells [5].

To confirm this hypothesis, we sorted pulmonary CD4^+^ T cells from naïve mice with the Tyto and compared their metabolic potential to cells sorted using beads. Pulmonary cells sorted on the Tyto resulted in generation of a CD4^+^ T cells pool exceeding 98% purity (data not shown), dramatically reducing contaminating cells from the lung preparations. We then examined the glycolytic rate of both cell populations. Irrespective of the purification method, total CD4^+^ T cells from naïve mice increased glycolysis after anti-CD3/CD28 bead stimulation (figure 3A). However, in comparison to bead sorted cells, basal ECAR levels were significantly reduced among Tyto sorted CD4^+^ T cells (figure 3B). Moreover, the basal ECAR observed among Tyto sorted cells was similar to that observed in immune populations in our previous publication [5]. These data highlight the importance of having a highly purified cellular population for downstream assays and confirm our hypothesis that contaminating cells in our bead purified CD4^+^ T cell samples contribute significantly to basal ECAR.

**Figure 3.**
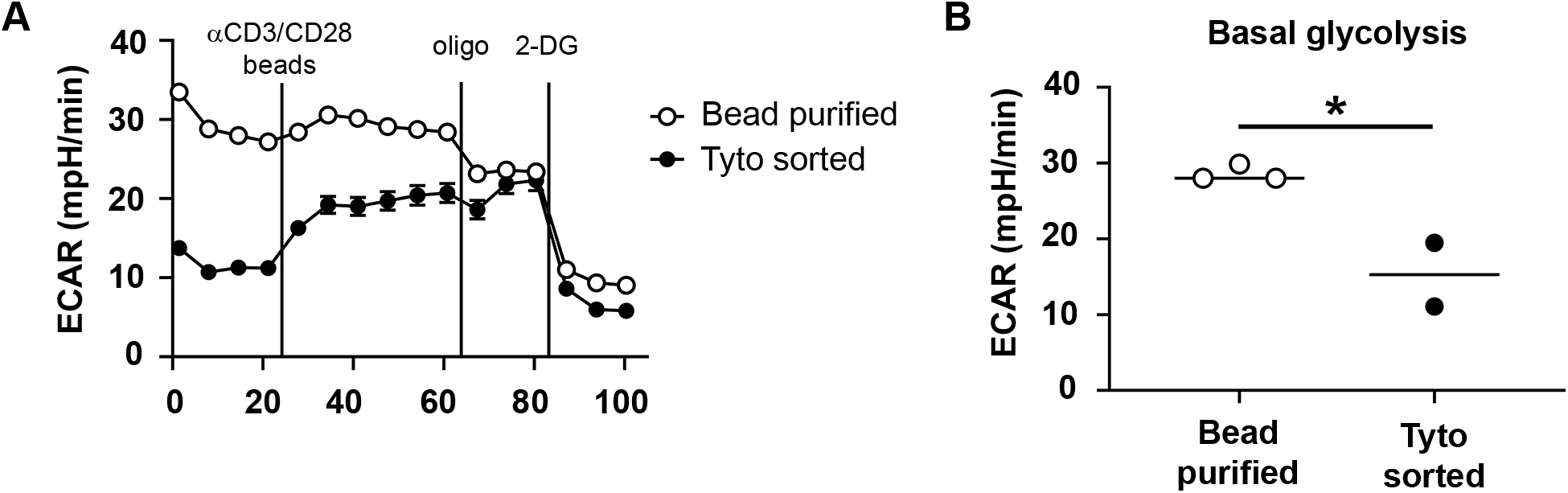
Contaminating cells increase basal glycolysis. Total CD4^+^ T cells were purified from naïve mice using CD4 beads or the Tyto and then seeded into a 96 well Seahorse plate. Cells were pooled from 5-10 mice per group with three to six technical replicates per group. A) The extracellular acidification rate (ECAR) was measured after activation in real-time with anti-CD3/CD28 beads and subsequent injection of oligomycin and 2-DG. B) The average basal glycolysis rate was determined. Panel A is representative of 2-3 independent experiments and panel B is pooled from 2-3 independent experiments. Error bars represent SEM. * *p* ≤ 0.05.

## Discussion

Isolation and downstream analysis of specific immune cell populations has been a critical component of immunological research over the last several decades. Over this same period, droplet-based sorters have constantly evolved to include additional lasers and/or channels, thereby allowing end-users to sort on an increasing number of fluorescent parameters. Although the sorting technology has improved with time, one feature remained consistent. That is, the danger of generating aerosols with infectious potential during the sort procedure. This small, but important feature, typically precludes the use of droplet-based sorters within containment laboratories. Some groups have circumvented this hurdle by placing the instrument in custom built, expensive, and large BSCs. However, given their expense and footprint, this is often not an option for most institutes working with BSL-3 and BSL-4 pathogens. To circumvent this challenge within our own BSL-3 laboratory, we acquired a MACSQuant Tyto (Tyto).

The primary feature of the Tyto that facilitates its bench top use in a containment setting is the ability to gently and rapidly sort cells within a sterile closed cartridge without generating aerosols. The Tyto uses microfluidics and extremely low air pressure to flow cells past the instrument’s lasers. A rapidly moving gate opens to divert a desired cell into the sort fraction and then closes again. This technology doesn’t rely on cells to be within droplets and they aren’t subjected to any charge changes to deflect them into a sort tube like traditional droplet-based sorters. Thus, this technology is not only appropriate for containment sorts, but is also attractive for use at BSL-1/BSL-2 for cell types that are extremely fragile and susceptive to mechanical perturbations. Although the Tyto may be appropriate for a variety of sorts outside and inside of containment, it should be emphasized that this instrument sorts one population at a time within the closed cartridge. Thus, indexed single cell sorts, sorting into plates, and sorting more than one population at a time are not possible.

Prior to implementation of the Tyto as a bench top sorter in the BSL-3, it was necessary to confirm that aerosols consistent with those capable of carrying infectious organism were not generated during operation of the equipment. We utilized a recently published and validated method for testing aerosol generation by droplet-based cell sorters to determine the aerosol generating potential of the Tyto [6]. This testing protocol was straightforward to complete and not cost-prohibitive. We utilized a commercially available vacuum, impactor cassette, and 1.0 μm internally fluorescent beads to determine whether the Tyto generated aerosols during the sort procedure. As a positive control, the in-house FACSAria II was placed in fail mode and clearly visible aerosols were collected using the same set-up. Importantly, no aerosols were generated by the Tyto during a sort of fluorescent beads. Overall, the testing procedure took less than 4 hours to collect and analyze the samples. Thus, it should be easily implemented by other institutions or groups to test their own instruments using the protocol outlined herein. The Tyto also has additional safety features built-in that are important to note during BSL-3 risk assessments including the inability to sort if there is a power failure, vacuum failure, an open door, or improper seating of the cartridge.

Once the equipment was validated for safety purposes, we next determined its experimental capacity. Since we do not possess a droplet-based sorter in our BSL-3 laboratory, there have been a number of research questions that were previously unanswerable because we lacked the necessary technology. We have implemented bead-based purification techniques for less complicated studies. However, when purifying cells from the pulmonary compartment these techniques result in approximately 85-90% pure cell populations, at best [5]. In some instances, this degree of purity is sufficient but in other experimental conditions, contaminating cells affect downstream analyses. For example, in a metabolic analysis of pulmonary CD4^+^ T cells, we were unable to sort cells from naïve mouse lungs due to time constraints. Instead, we used a bead-based purification scheme, which resulted in CD4^+^ T cells that were approximately 85% pure. Although these contaminating cells were not affected by our anti-CD3/CD28 stimulation, they were metabolically active and thus artificially elevated the basal glycolysis measurement of the sample. As a consequence, it appeared as though naïve CD4^+^ T cells had higher basal glycolysis rates compared to CD4^+^ T cells from immune animals isolated via sorting. The ability to sort much faster on the Tyto allowed T cells to be sorted from both naïve and immune animals prior to downstream analysis. As predicted, the near elimination of contaminating cells from our naïve CD4^+^ pool reduced their basal glycolysis to levels comparable to immune CD4^+^ T cells (figure 3).

As noted above, sorting cells on the Tyto saved a significant amount of time, which is critical for cell viability and the ability to perform accurate downstream analysis. Time is also saved outside of the actual sort. While both traditional droplet-based sorters and the Tyto need time for the lasers to warm-up, the Tyto does not require the operator to set-up and optimize the stream. Once the sort is completed, the Tyto can simply be shut down. There is no decontamination procedure to clean the fluidics lines in preparation for the next user, nor is there waste fluid to dispose of like one must deal with for droplet-based sorters. The only waste generated from the sort is the cartridge itself which can be disposed of in the typical biohazard waste stream. The closed cartridge system not only eliminates some set-up and shut-down time, it also prevents sample loss. For example, if it is determined that the number of sorted cells was insufficient for downstream analysis, the user can return to the cartridge’s negative fraction and sort the desired cell population a second time. This is not possible on droplet-based sort systems because all unwanted cells are channeled into the waste. Together, Tyto sorts are more efficient with no additional set-up or shut down time, minimal to no cell loss, and does not generate large amounts of liquid waste requiring appropriate decontamination prior to its disposal.

While the Tyto has many advantages compared to droplet-based sorters, one caveat is that use of this technology requires the user to design antibody panels differently. Unlike traditional droplet-based sorters where the same analysis antibody panel could be used to sort cells, the Tyto has strict requirements that must be met during panel design. Specifically, the desired cell population must be positively stained with 2 fluorochrome-conjugated antibodies that are excited by 2 different lasers. While this can be challenging, there are often two densely expressed markers on a cell population of interest. For example, we have used Thy1 as a common T cell marker and CD45 for macrophage sorts. Another important consideration for use of the Tyto is compensation. To date, our sorts have not utilized more than 4 fluorochrome-conjugated antibodies on a single cell population of interest. We have been successful in designing panels with to minimize fluorophore bleed-through to other channels. However, one potential downside with using the Tyto to sort a cell population that requires numerous markers for positive identification is the necessity to run compensation controls. Unlike a droplet-based sorter where the compensation controls could be prepared in tubes and those quickly run to establish the compensation matrix, running compensation on the Tyto would be more cumbersome having to switch the compensation control within a cartridge. We recommend Tyto users run pilot sorts to identify optimal antibody panels and sort parameters to maximize target cell recovery.

In summary, the ability to sort immune cell populations in high containment (BSL-3/BSL-4) laboratories has historically been hampered by risk of aerosol generation by droplet-based cell sorters. While these risks can be mitigated by engineering controls such as placing the instrument in a BSC, these are often cost- or space-prohibitive. As an alternative, we have utilized a Tyto cell sorter in our BSL-3 laboratory. This instrument not only has a small footprint, but most importantly does not generate aerosols during the sort procedure. The use of this technology will uplift current immunological research in containment laboratories by allowing a greater number of research groups to isolate cell types involved in the immune response to high consequence pathogens for down-stream applications.

